# Single-Cell Dissection of Immunometabolic Rewiring in the Porcine Ileum during *Salmonella* Typhimurium Infection

**DOI:** 10.64898/2025.12.04.690681

**Authors:** José M. Suárez-Cárdenas, Tomàs Montserrat-Ayuso, Mónica Alfonso-Núñez, Anna Esteve-Codina, Raúl Fernández-Rodríguez, Antonio Romero-Guillén, Tránsito García-García, Cristina Arce, Amparo Martínez Martínez, Juan J. Garrido, Sara Zaldívar-López

**Author notes:** **Corresponding author**: Sara Zaldívar-López. These authors share senior authorship (Juan J. Garrido, Sara Zaldívar-López).

## Abstract

*Salmonella* Typhimurium is a major zoonotic pathogen, with pigs acting as important subclinical carriers. To explore the specific intestinal immune response at the cellular level, we used Single-cell RNA sequencing (scRNA-seq), enabling detailed analysis of immune cell types and gene expression profiles during infection. In addition to enterocytes, our results revealed the presence of diverse immune populations, including monocytes/macrophages, dendritic cells, innate lymphoid cells (ILCs), thirteen T cell subtypes and five B cell populations were identified, revealing pronounced infection-driven alterations in cellular composition and transcriptional states. Among T cells, naïve and follicular CD4^+^/CD8^+^ αβ T cells and NK T cells were expanded, whereas effector CD8^+^ T cells and CD2^−^ and SELL^hi^ γδ T cells were decreased. B-cell populations shifted toward activated and cycling states, with decreased antibody-secreting, resting, and transitioning cells. Dendritic cells and monocyte/macrophage populations were expanded, and group 3 ILCs and enterocytes were markedly reduced. Pathway analyses revealed robust cell type-specific immunometabolic remodeling, including enhanced protein-folding and stress-adaptive pathways in T and B cells, heightened inflammatory, interferon, and cytokine signaling in myeloid populations, and coordinated metabolic and immune adjustments in epithelial cells, highlighting the complexity of host responses to *Salmonella* infection. This study provides the first scRNA-seq landscape of the porcine ileum during *S.* Typhimurium infection, offering insight into host immune cell dynamics and immunometabolic responses.

## INTRODUCTION

Salmonellosis, caused by bacteria of the genus *Salmonella*, remains one of the most prevalent foodborne zoonoses worldwide, representing major public health and veterinary concerns. Non-typhoidal *Salmonella* (NTS), particularly *Salmonella enterica* subsp. *enterica* serovar Typhimurium (hereinafter *S*. Typhimurium) and *Salmonella enterica* subsp. *enterica* serovar Enteriditis, are leading causes of gastroenteritis in humans, with global estimates suggesting tens of millions of infections each year. In 2023, the European Union (EU) reported 77,486 confirmed cases of human salmonellosis (18 cases per 100,000 population), which shows a 16.9% increase compared with 2022 (15.4 cases per 100,000 population), and resulting in 88 reported deaths (1). Salmonellosis remains the second most frequently reported zoonosis in the EU, and pigs and pork products constitute a major reservoir and source of human infection (2). In humans, the disease typically causes diarrhea, abdominal pain, nausea and vomiting, and although it is usually self-limiting in healthy individuals, it can become severe or even fatal in infants, elderly people or immunocompromised patients (3). Beyond its public health impact, salmonellosis also imposes substantial economic losses on livestock production due to reduced productivity, trade restrictions, and control costs. In pigs, infection may manifest as acute gastroenteritis in young animals, and they can also remain as subclinical or asymptomatic carriers. The bacteria persist in intestinal tissues or mesenteric lymph nodes and being intermittently shed into the environment (2, 4, 5). Consequently, controlling *Salmonella* infections in pigs is critical not only for animal welfare and productivity but also for mitigating the risk of zoonotic transmission to humans. Given the global and European burden of salmonellosis, a One Health approach integrating human, animal and environmental perspectives is essential.

Upon oral ingestion of contaminated food or water, *S*. Typhimurium passes through the gastrointestinal tract and reaches the small intestine, where it preferentially colonizes the ileum, interacting with the intestinal epithelium and the resident microbiota, constituting the first line of defense. Here, the host will trigger an initial immune response, aided by local specialized lymphoid structures such as Peyeŕs patches within the gut-associated lymphoid tissue (GALT), that will orchestrate adaptive immune response by supporting B and T cell activation (6, 7). For the attack and colonization, *S*. Typhimurium employs its virulence machinery as type III secretion system (encoded by *Salmonella* Pathogenicity Island -SPI-1) to inject effector proteins into epithelial cells, triggering cytoskeletal rearrangements and facilitating bacterial uptake and internalization via macropinocytosis (8, 9). Once internalized, the bacteria reside within membrane-bound *Salmonella*-containing vacuoles (SCVs), where SPI-2 mediates intracellular survival and replication by subverting host antimicrobial pathways, particularly in phagocytic cell such as macrophages and dendritic cells (10, 11). The host immune system rapidly recognizes the pathogen via pattern recognition receptors, initiating an innate immune response characterized by recruitment of neutrophils, monocytes/macrophages and natural-killer (NK) cells to the site of infection (12). The resulting local inflammatory response produces antimicrobial molecules and immune mediators while simultaneously modifying the tissue microenvironment, which can be exploited by the pathogen to persist and replicate within SCVs. Subsequently, adaptive immune responses, including T and B cell activation and antibody production, are engaged and play a crucial role in controlling infection and determining pathogen clearance versus persistence (13). The interplay between innate and adaptive immunity, together with the heterogeneity and transcriptional programs of these immune populations, critically shapes infection outcomes and bacterial persistence, highlighting the importance of dissecting these cellular networks at high resolution.

The ileum represents a complex immunological environment where multiple cell types interact to maintain tissue homeostasis and coordinate responses to invading pathogens. Within this site, epithelial cells, B and T lymphocytes, innate lymphoid cells (ILCs), macrophages, dendritic cells, and other immune populations form an intricate network that determines the outcome of *Salmonella* infection (14, 15). Understanding the heterogeneity, relative abundances, and gene expression profiles of these populations is essential for elucidating how local immune responses are orchestrated and how specific cell types may support bacterial persistence or clearance. To our knowledge, this study represents the first single-cell transcriptomic analysis of the porcine ileum during *Salmonella* infection, providing unprecedented insight into host cellular responses. Sc-RNA-seq has emerged as a powerful tool to dissect such complexity, enabling the identification of discrete cellular subsets, activation states, and transcriptional programs that would otherwise be masked in bulk tissue analyses (16, 17).

Recent advances in the field of immunometabolism have revealed a tight interconnection between cellular metabolic programs and immune cell function. The activation, differentiation, and effector potential of immune cells are profoundly influenced by their metabolic state, which regulates cytokine secretion, phagocytosis, antigen presentation, and overall antimicrobial defense (18, 19). Within the intestinal microenvironment, metabolic adaptation is particularly critical due to the high antigenic load, constant microbial exposure, and fluctuating nutrient availability. In the context of *Salmonella* infection, the pathogen itself actively reprograms host metabolic pathways to establish a favorable intracellular niche, while host immune cells undergo substantial metabolic reprogramming to sustain antimicrobial responses (20). Understanding these immunometabolic interactions is key to uncovering how infection outcomes are determined and how pathogens exploit host metabolism to persist. Considering previous evidence, here we hypothesize that *Salmonella* infection induces alterations in both the abundance and transcriptional programs of ileal immune and epithelial populations, with distinct biological consequences for immune functions and signaling pathways. This study aims to provide a comprehensive understanding of host cellular responses and uncover the mechanisms underpinning bacterial persistence and disease progression.

## RESULTS

To verify the efficiency of *Salmonella* infection in porcine ileum, the presence of bacteria in the tissue was determined by *Salmonella* DNA quantification in ileum content (using a *S*. Typhimurium-specific probe), and detection of *Salmonella* by immunohistochemistry and immunofluorescence using *Salmonella*-specific antibodies (**Figure 1**). We observed a significant increase in *Salmonella* DNA (as shown by decreased qPCR amplification Ct) when compared to control samples (**Figure 1A**). Similarly, in tissue sections of control animals we could observe an intact mucosa, preserving the intestinal villi and without the presence of bacteria (**Figures 1B, 1C**), meanwhile in infected animals intestinal villi were atrophied and epithelial damage could be observed, in addition to the presence of bacteria in the epithelial barrier and Peyeŕs patches (**Figures 1D, 1E, 1F**).

**Figure 1.**
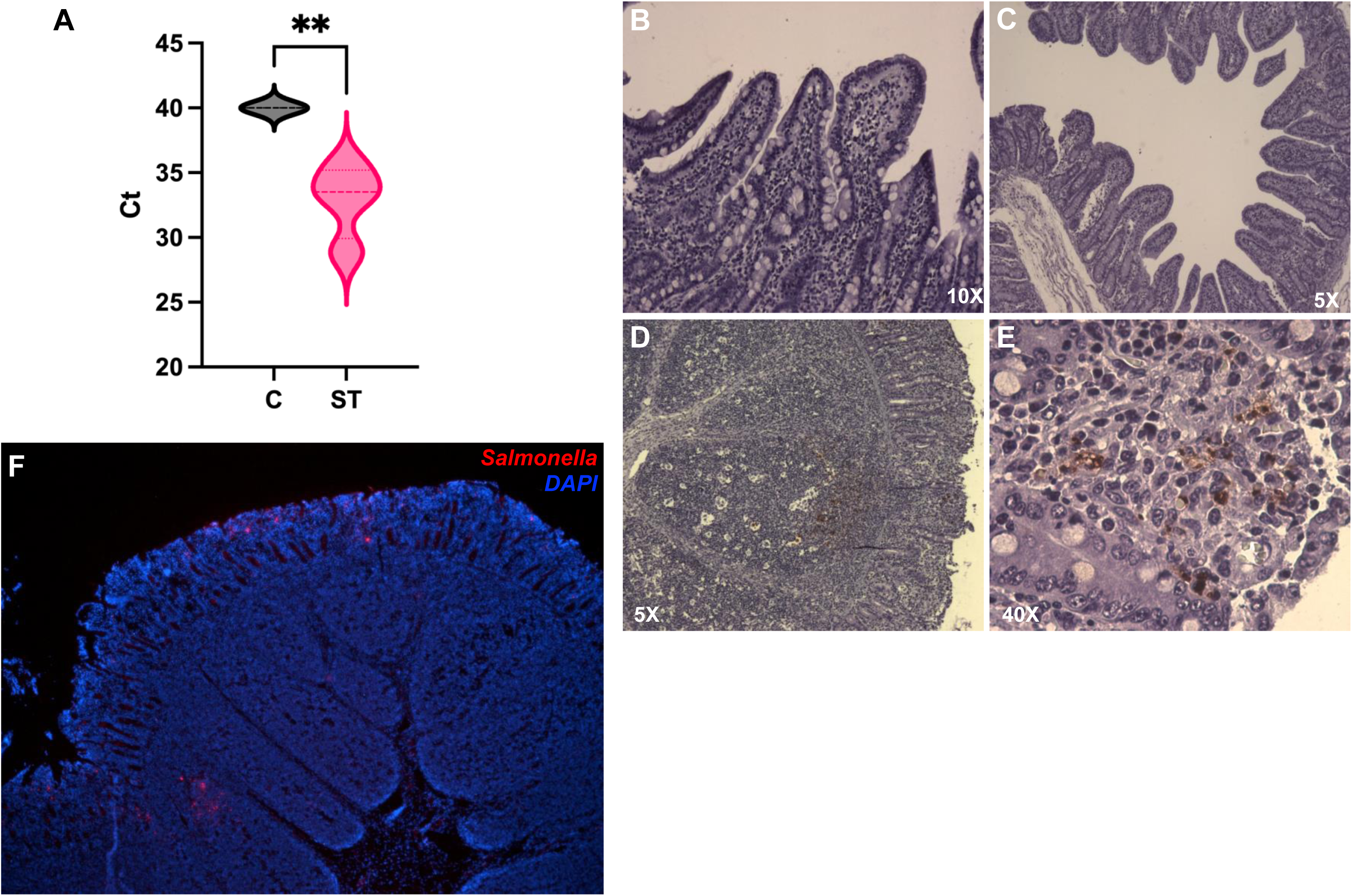
*Salmonella* Typhimurium DNA quantification in ileum fecal content (**A**). Immunohistochemical detection of *Salmonella* Typhimurium in porcine ileum sections of control animals (**B** and **C**) and infected animals (**D** and **E**) at 2 days after oral inoculation. *Salmonella* Typhimurium detection by immunofluorescence in ileum sections (**F**). **: p<0.01

### Cell population identification and annotation

Initial annotation included a high abundance of T and B cells, and myelomonocytic cells such as monocytes/macrophages and dendritic cells, along with enterocytes (**Figure 2**). A second level of analysis was performed to further characterize these populations, where cells were separated according to their lineage (ILC, B or T cells) to further determine the cellular subtypes of each group more precisely. Thus, two ILCs, thirteen T cells and five B cell subtypes were identified (**Figure 3**). Among the T and ILC cell subtypes annotated we found: Activated CD8^+^ αβ T cells, Naive CD4^+^/CD8^+^ αβ T cells, Cycling CD4^+^/CD8^+^ αβ T cells, Cytotoxic CD8^+^ αβ T cells, Follicular CD4^+^ αβ T cells, IFN-responsive CD8^+^ T cells, Naïve CD8^+^ T cells, NK CD8^−^,NK CD8^+^,Non-naïve CD4^+^ αβ T cells, Non-naïve CD8^+^ αβ T cells,CD2^−^ γδ T cells, SELL^hi^ γδ T cells, Activated group 1 ILCs, and Group 3 ILCs (**Figure 3**). Among the B cell subtypes we found Cycling B cells, Resting B cells, Antibody-secreting cells, Transitioning B cells and Activated B cells (**Figure 4**).

**Figure 2.**
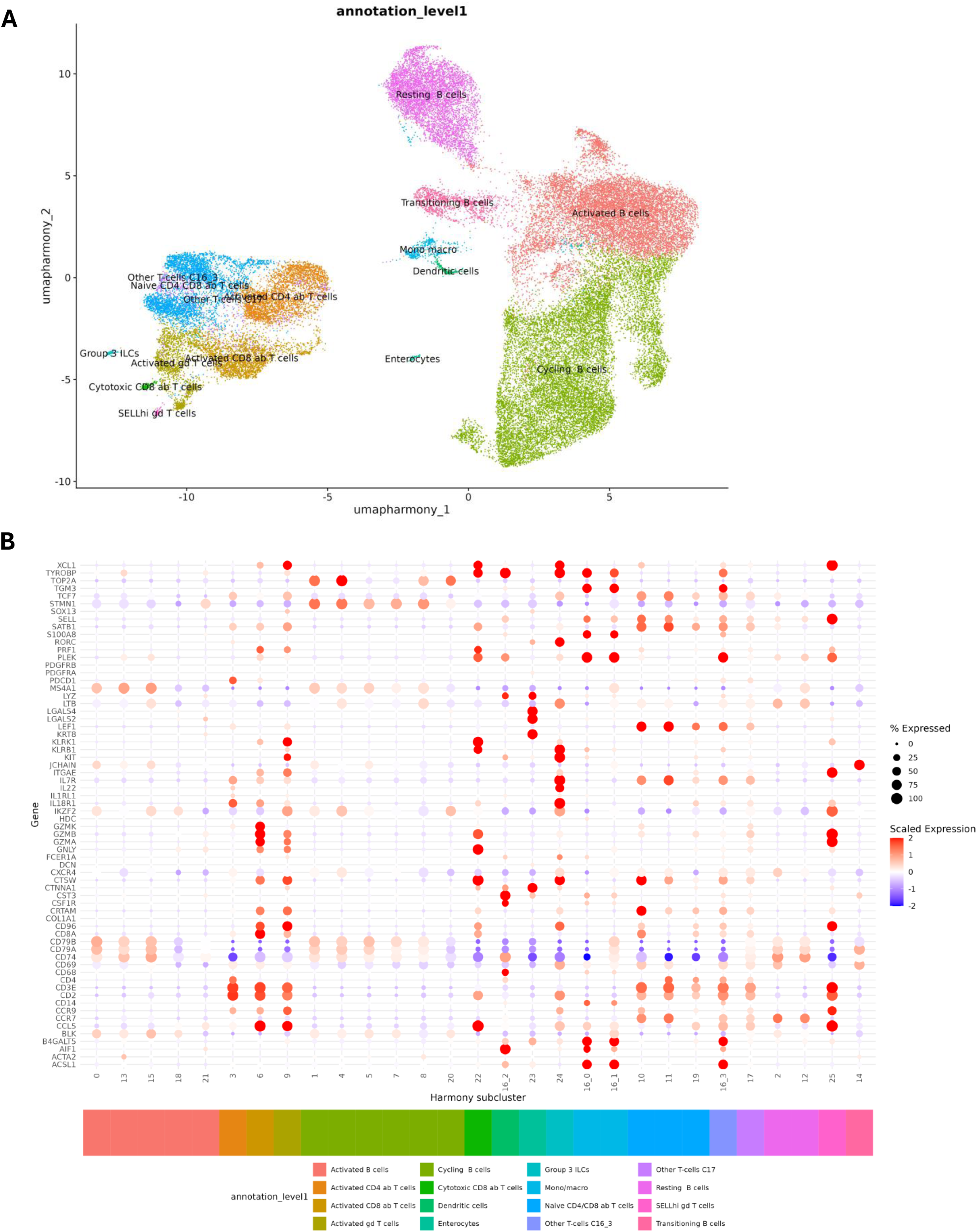
Definitive annotation of the ileum cells (**A**). Annotation was performed based on marker expression of cells; selected cell markers are selected in (**B**), showing expression in each cell type.

**Figure 3.**
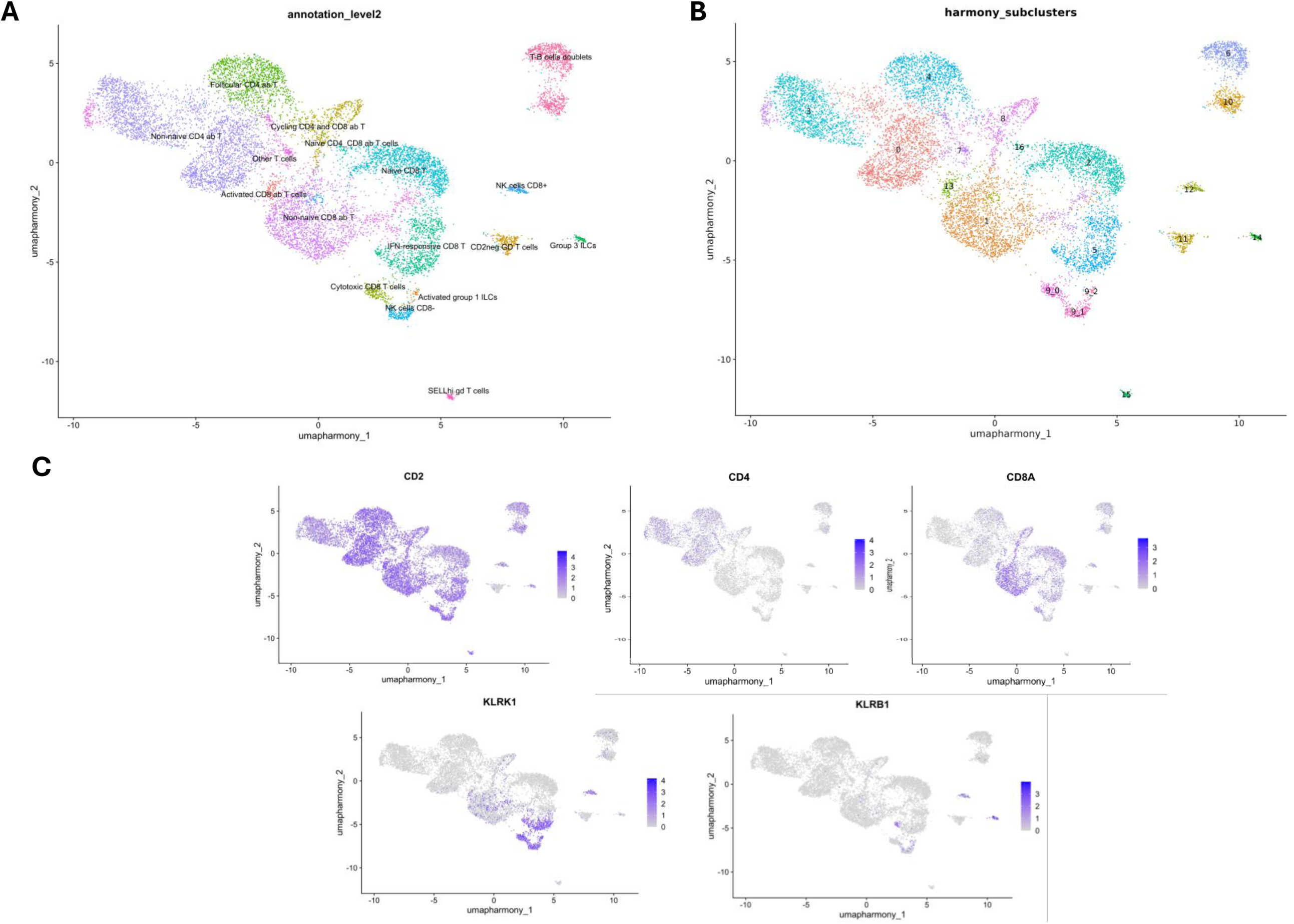
Annotation of T cell populations (**A**), based on initial clustering shown in (**B**). Cell populations were identified by expression of population-specific gene markers (**C**) such as CD2, marker of T cells; CD4 and CD8A, are markers of the CD4^+^ and CD8^+^ T cell subtypes; and expression of KLRB1 and KLRK1, markers of NK cells.

**Figure 4.**
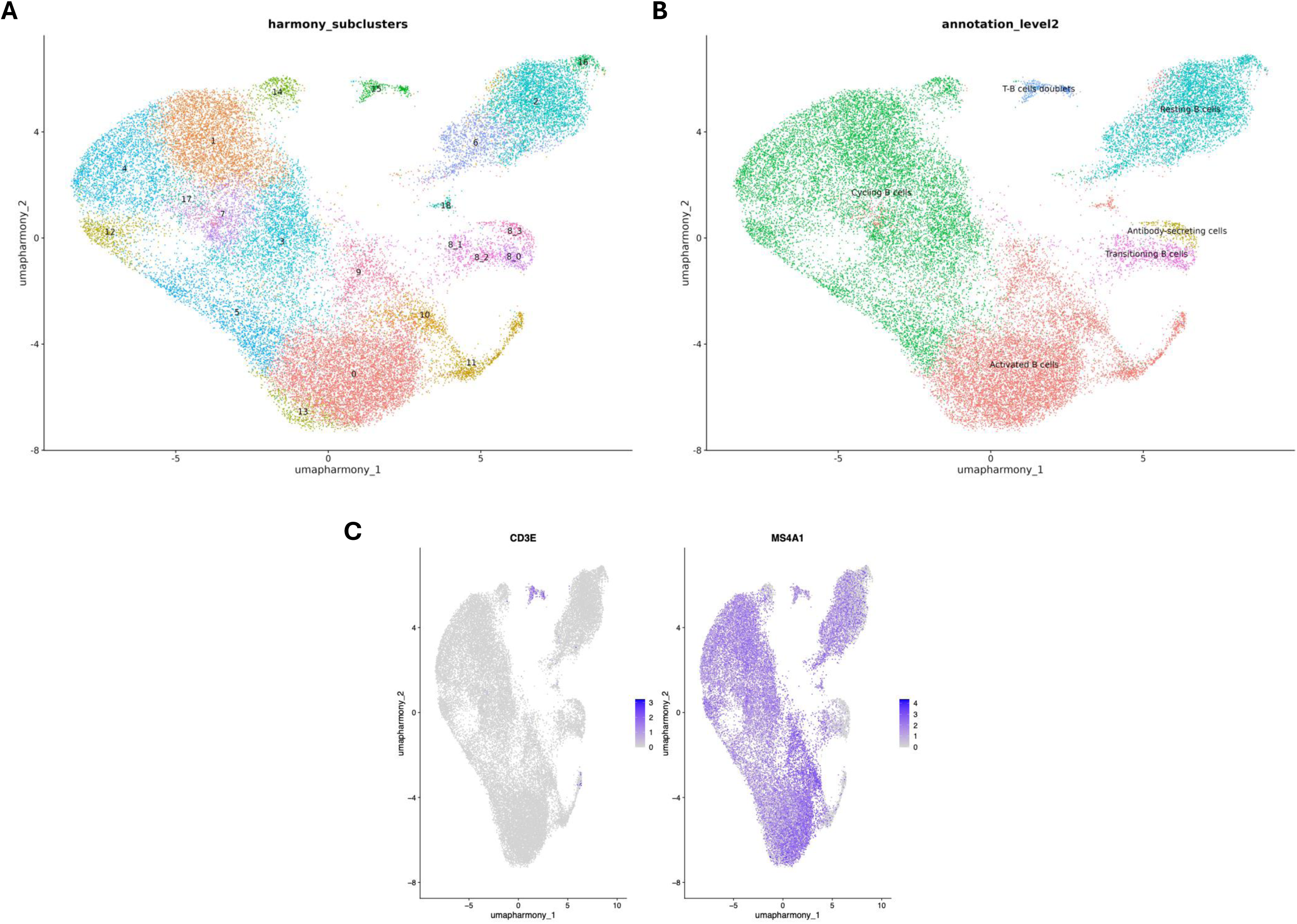
Annotation of B cell populations (**A**), based on initial clustering shown in (**B**). Cell populations were identified by expression of population-specific gene markers such as MS4A1, and negative to T cell marker CD3E (**C**).

### Changes in cell type distribution during S. Typhimurium infection in porcine ileum

Once cells were properly annotated, a comparative study of the abundance of the different cell populations was performed between control and infected animals, to determine whether *Salmonella* induces changes in the number of cellular subtypes. The number of cells from all samples of each condition belonging to each cell subtype was analyzed, revealing significant alterations in the abundance of several immune cell populations in the ileum of infected pigs when compared to uninfected controls. In general, we observed a high abundance of B cells in porcine ileum, with T cells being also highly abundant in this tissue (**Figure 5**). At first level annotation (**Figures 5A, 5B**), B cells showed a heterogeneous response: activated B cells were significantly increased in infected animals, meanwhile cycling B cells, resting B cells, and transitioning B cells were significantly reduced in this group. T cell populations displayed diverse patterns: naïve CD4^+^ and CD8^+^ αβ T cells, as well as cytotoxic CD8^+^ αβ T cells, were significantly increased, while activated CD8^+^ αβ T cells, activated γδ T cells, and SELL^hi^ γδ T cells were significantly decreased. Additionally, we observed that dendritic cells and monocyte/macrophage populations were significantly expanded in ileum from infected animals, whereas abundance of group 3 innate lymphoid cells (ILCs) and enterocytes was significantly reduced.

**Figure 5.**
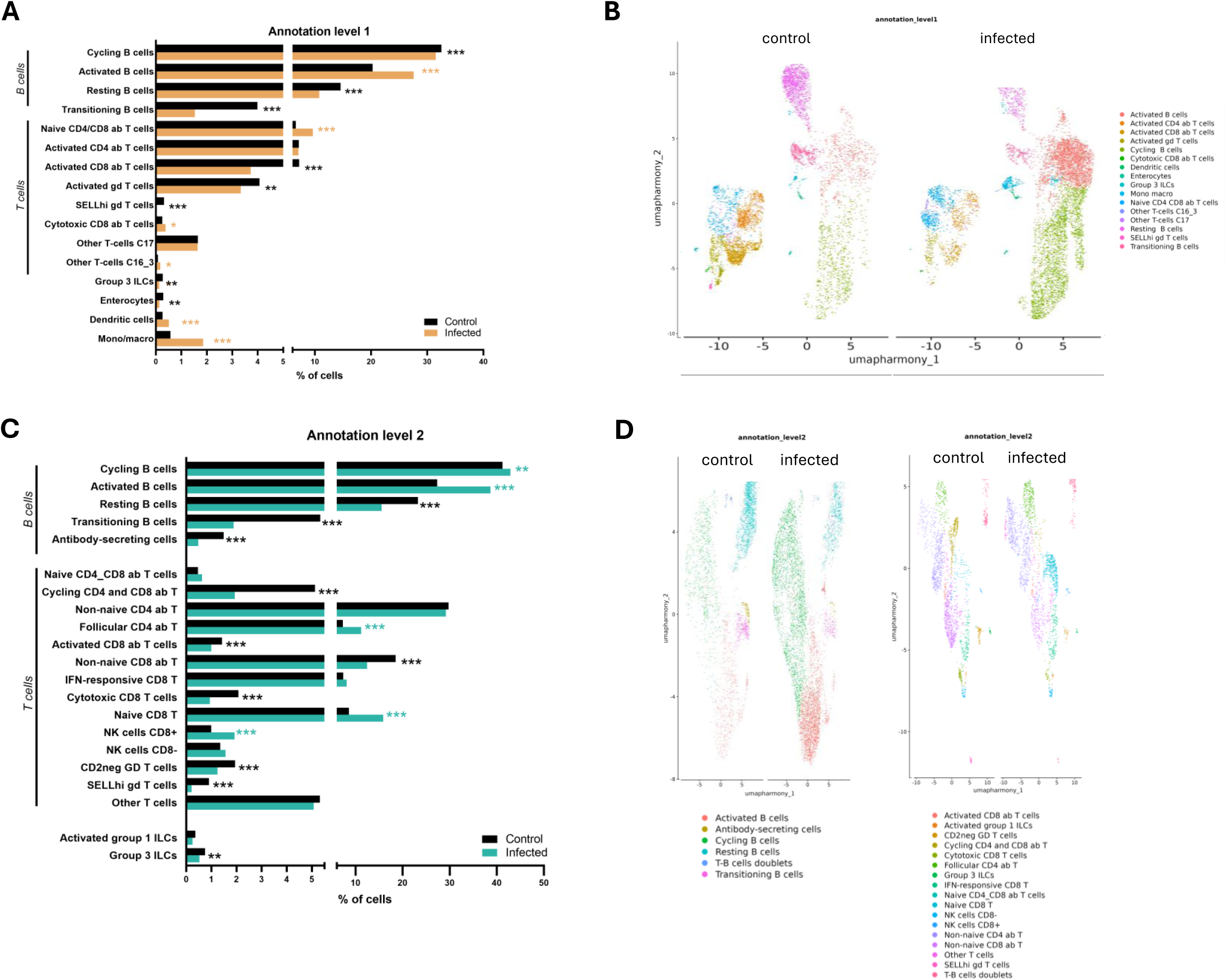
Analysis of abundance (% of all cells) of different cell populations found in the porcine ileum and comparison between infected cells and controls at a broader and more general level (**A**) and representative umap graph of control and infected individuals (**B**). Detailed analysis of abundance (% of all cells) of T and B cell subsets in the porcine ileum and comparison between infected cells and controls (**C**), and representative umap graph of control and infected individuals (**D**). *: p<0.05; **: p<0.01; ***: p<0.001; ****: p<0.0001.

A more detailed annotation level (level 2, **Figures 5C, 5D**), allowed the identification of further alterations in infected animals compared to controls. B cell subpopulations displayed a significant increase in activated and cycling B cells, whereas antibody-secreting, resting, and transitioning B cells were significantly decreased in infected animals. T cell analysis showed an increase in follicular CD4^+^ αβ, naive CD8^+^, and NK T cells. On the other hand, cycling CD4^+^/CD8^+^, activated, cytotoxic and non-naïve CD8^+^, as well as CD2^−^ and SELL^hi^ γδ T cells were significantly reduced in infected ileum compared to uninfected controls. Additionally, the number of group 3 ILC cells was reduced in ileum from infected animals. Naïve CD4^+^/CD8^+^ αβ, IFN-responsive CD8^+^, and NK CD8^−^ T cells also showed increased abundance in infected ileum, but changes were not statistically significant.

### Gene expression profiling of distinct cell populations and impact on biological functions

For each cell population identified, a differential gene expression analysis was conducted to investigate the effects of *Salmonella* infection, and the immunometabolic pathways that may be subsequently altered. Given the low number of significantly dysregulated genes in some populations, we lowered the threshold to uncorrected *p value* <0.05 and absolute fold change >2 to further understand the biological processes altered by the gene expression changes (**Supplementary file 1**). As we can see in **Figure 6D**, the T-cell subtypes with the highest number of differentially expressed genes are non-naïve CD4^+^, follicular CD4^+^ and IFN-responsive CD8^+^ cells. From the B cell lineage, cycling, activated and resting B cells have a high number of differentially expressed genes during infection, with cycling cells showing predominant upregulation and activated cells indicating a profound downregulation.

**Figure 6.**
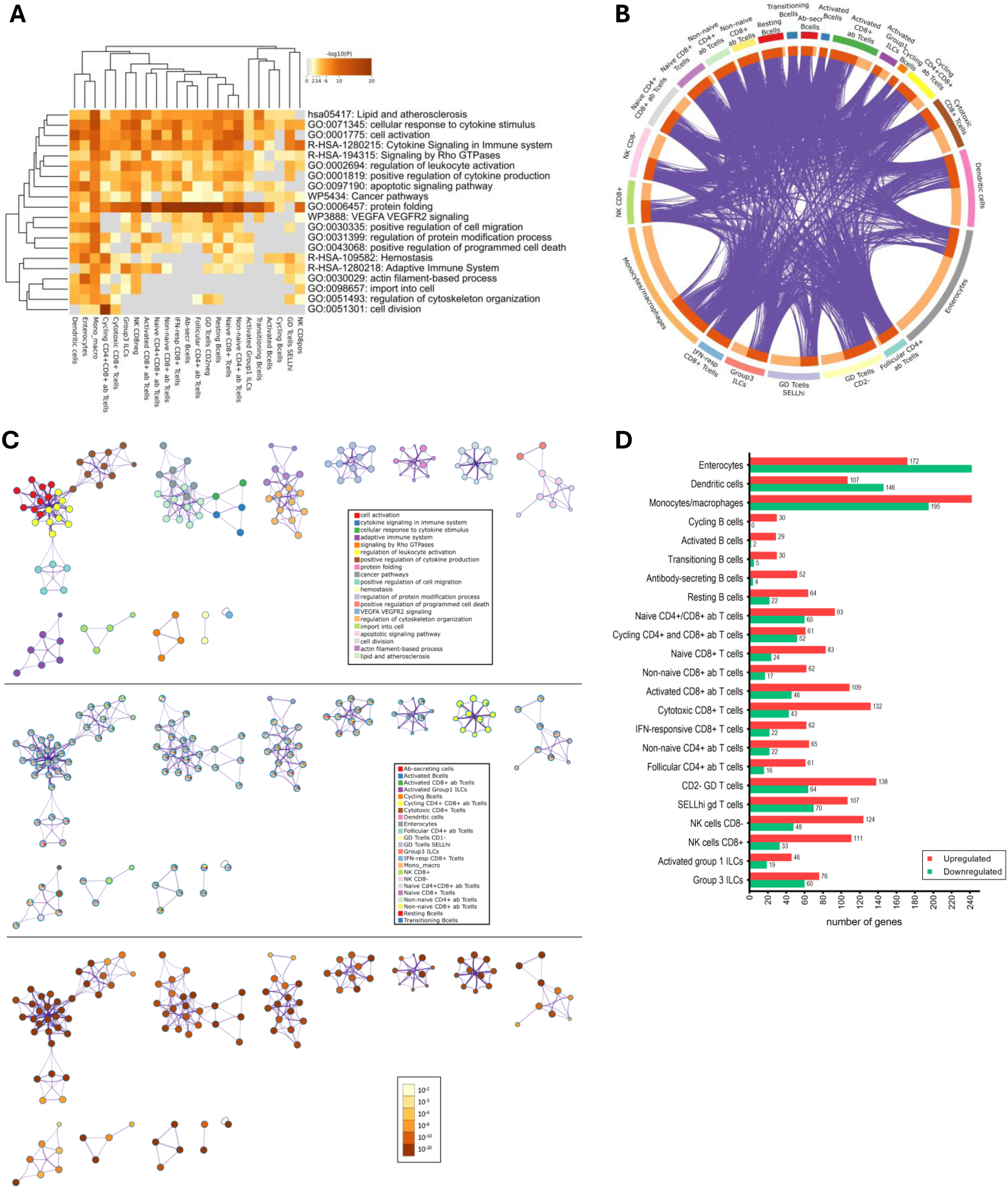
Functional enrichment analysis revealed enriched ontologies and terms across all cell types (colored by p-values) (**A**). Several genes in the different cell subtypes were shared (purple lines link identical genes; genes that hit multiple cell types are dark orange, and genes unique to a list are shown in light orange) (**B**). Network of enriched terms: nodes colored by cluster ID (upper image); nodes colored as pie charts of different cell subtypes (intermediate image); nodes colored by p-value (lower image) (**C**). Differentially expressed genes (p value < 0.05 and |FC|=2) in the immune cell populations identified in porcine ileum infected with *Salmonella* Typhimurium (**D**).

All cell populations were analyzed using Metascape tool (21). The analysis showed many differentially expressed genes that were shared between the different cell populations (**Figure 6B**), and the top 20 enriched terms (pathways and functions) across the cell populations studied (**Figure 6A**). Among these functions we observe that dysregulation of immune functions such as cytokine signaling, leukocyte activation, and adaptive immunity are present in most cell populations from ileum, as well as metabolic processes (lipid signaling) and transport into cell. Cell migration is affected mainly in monomyelocytic cells (monocytes/macrophages and dendritic cells). Alteration of cytoskeletal reorganization, Rho GTPases signaling and actin filament processes is present in enterocytes and monocytes/macrophages. To better illustrate how these terms relate to one another, a subset of enriched terms were selected and displayed as a network graph. Terms were clustered by function, cell types where each term was dysregulated is shown by colors within the term, and p-values are also indicated in **Figure 6C**. Here we observed that some functions are altered in many of the cell types in ileum (e.g., leukocyte activation and immune responses), and other seem to be altered only in specific cell types: cell division is specific of cycling CD4^+^ and CD8^+^ αβ T cells; import into cell is altered in NK CD8^+^ cells; and dysregulation of actin filament-based process is present in cytotoxic CD8^+^ and NK CD8^+^ cells.

Gene set enrichment analysis (GSEA) results revealed cell type-specific patterns of pathway activation and suppression, highlighting heterogeneous response to *Salmonella* infection (**Supplementary file 2**). Comparisons between infected and control samples allowed the identification of pathways that were commonly affected, as well as those selectively modulated in specific cellular lineages. Activation of protein maturation, chaperone-mediated protein folding, and general protein folding pathways were found in several cell types, along with responses to temperature stimulus (**Figure 6A**). In particular, transcriptomic signatures indicative of inflammatory, immunomodulatory, and stress-related responses were observed in both epithelial and immune cell populations. The following sections describe expression changes in each cell type, emphasizing those with the greatest functional relevance in the context of *Salmonella* infection.

#### Enterocytes

Intestinal epithelial cells respond to *S.* Typhimurium infection by adjusting both metabolic and immune programs. In response to bacterial infection, enterocytes displayed a coordinated downregulation of metabolic pathways involved in amino acid (NES= −1.9) and lipid metabolism (NES= −1.6) as indicated by subexpression of *PPARG*, *GPX4*, *HNF4A*, and *LPL*, suggesting a reduction in nutrient processing and energy generation under infection conditions. In contrast, multiple immune-related pathways were upregulated, including humoral immune responses (NES= 2.02, by upregulation of *CXCL2, CXCR4*, *C1S*), and cellular responses to abiotic and environmental stimuli (upregulation of *TIMP1*, *CD40*, *IRF1*). Additional enrichment was observed in ribonucleoprotein complex biogenesis, positive regulation of monoatomic ion transport, and processes associated with cell–cell fusion and syncytium formation. *TIMP1* is also involved in integrin mediated signaling.

#### Monocytes/Macrophages

Functional enrichment analysis of differentially expressed genes revealed activation of multiple pathways related to immune response, metabolism, and cellular stress. Inflammatory pathways such as inflammatory response (NES= 2.07), response to molecules of bacterial origin (NES= 2.03), response to lipopolysaccharide (NES= 1.98), cytokine production (NES= 1.56), and type I interferon response (NES= 1.91) were upregulated, consistent with the strong induction of *CXCL8*, *NFKBIA*, *IRF1*, *IRF7*, *NLRP3* and *GBP1*, reflecting a robust activation of NF-kB and interferon signaling. Concurrently, pathways related to immune regulation, including negative regulation of T cell activation and proliferation (NES= 1.93, 1.83), and negative regulation of leukocyte cell-cell adhesion (NES= 1.93), driven by upregulation of *CD274* (PD-L1) suggest potential immunomodulatory effects on T cell responses. Metabolic pathways such as glycolytic process (NES= 1.90), pyruvate metabolic process (NES= 1.91), and amino acid catabolic process (NES= 1.65) were also enriched, in agreement with the upregulation of *HK3*, *PKM*, *IDO1*, *ACOD1,* and *PGAM1*. Additionally, increased expression of *FTL*, *SOD2*, and *HSPA6* points to enhanced oxidative and protein stress responses, as well as iron sequestration and lipid metabolism remodeling. Downregulation of *EPAS1* and *LPIN1* points to reduced lipid biosynthesis and hypoxia signaling. Finally, activation of cellular stress and turnover pathways, including protein folding, protein maturation, cellular catabolic process, apoptotic process, and programmed cell death, indicate heightened cellular stress and protein homeostasis challenges in monocytes/macrophages during infection. In addition, several downregulated genes (*ITGA2*, *PLXNC1*, *RAP1GAP2*, *RAP2A*, *DOCK1*, *GSN*, and *CYTIP*) suggest repression of pathways controlling cytoskeletal remodeling, cell adhesion, and migration, while *TGFBR2*, *JAML*, and *PALS2* may reflect altered intercellular communication and antigen presentation, and lower expression of *EEF2K* and *MAST2* suggests diminished protein synthesis and stress adaptation capacity.

#### Dendritic cells

Although no significant functions were retrieved, upregulated genes reflected activation of cellular and oxidative stress pathways (*HMOX1*, *HSPA6*, *BAG3*), inflammatory and chemotactic responses (*SAA3*, *IRF1*, *CCL3L1*), and interferon-mediated antimicrobial defense (*IRF1*, *FTH1*). Conversely, several genes linked to dendritic cell maturation, migration, and antigen presentation (*CCR7*, *SLAMF9*, *GPR183*, *SAMSN1*, *FCER1A*, *MS4A2*) as well as amino acid and glycosylation metabolism (*ST6GALNAC3*) were downregulated, suggesting impaired T cell communication and altered metabolic function.

#### Innate lymphoid cells (ILCs)

Group 3 ILCs show an activation of immune response by upregulation of TNF signaling genes (*CSF*, *IRF1*, *TRAF1*, *NKBIA*) and MAPK signaling (*MAP3K5*, *MAP3K8*, *KIT*). Group 1 ILCs show upregulation of antigen presenting genes, both MHC class II (*SLA-DRB1*, *CD74*, *SLA-DQB1*) and MHC class I (*SLA-2*, *PDIA3*). Regulation of T cell activation (*HSPD1*, *IRF1*) and interferon gamma response (*IRF1*, *GBP1*, *XCL1*).

#### Activated B cells

Activated B cells increase both migration and biosynthetic activity in response to infection. In infected animals, although mild activation of inflammatory genes is observed (*NFKBIA*), activated B cells exhibited a pronounced downregulation of chemotaxis-related pathways (NES= −1.88), indicating decreased mobilization within the infected tissue. Similarly, pathways associated with peptide biosynthesis (NES= −1.46), and amide metabolism (NES= −1.46), and translation were downregulated, reflecting a decreased metabolic and biosynthetic activity to support cellular activation.

#### Antibody secreting B cells

Functional enrichment analysis of antibody-secreting cells revealed an increase in aerobic respiration (NES= 1.74) and oxidative phosphorylation (NES= 1.74) by upregulation of *NDUFS8*, indicating a predominant reliance on oxidative metabolism to sustain their biosynthetic demands. Genes related to antigen presentation in class I and II are also upregulated, such as *PDIA3*, *SLA-2* and *SLA-DRB1*.

#### Cycling B cells

In infected animals, cycling B cells showed a pronounced downregulation of multiple biosynthetic and translational pathways, including peptide and amide biosynthesis (NES= −1.6), rRNA processing (NES= −1.71), ribosome biogenesis (NES= - 1.97), and cytoplasmic translation (NES= −1.98). This suggests a temporary reduction in global protein production, which may reflect a strategic allocation of resources during rapid proliferation. There is an upregulation of genes of innate (*NFKBIA*), interferon response (*IRF1*) and regulation of protein ubiquitination (*CDC20*).

#### Resting B cells

Resting B cells exhibited upregulation of pathways related to mitochondrion organization (NES= 1.62, *NDUFS8* upregulation). In contrast, pathways involved in peptide and amide biosynthesis (NES= −1.62, −1.53) and translation (NES=-1.61), were downregulated, indicating a reduced demand for global protein synthesis under resting conditions.

#### Transitioning B cells

Transitioning B cells exhibit coordinated activation of defense and protein handling pathways, reflecting their shift from a resting to a more functionally responsive state. In infected animals, transitioning B cells showed strong upregulation of pathways related to defense response (NES= 1.64), innate immune activation (NES= 1.71), and responses to biotic stimuli (NES=1.70), including responses to viruses and other organisms, as shown by upregulation of *CXCL10*, *NFKBIA and IRF1*. In contrast, pathways associated with cytoplasmic translation, general translation, and peptide and amide biosynthesis and metabolism were downregulated (NES= −1.86 to −1.75), suggesting a reduction in global protein synthesis during this transitional phase.

#### Activated CD8^+^ αβ T cells

During *S.* Typhimurium infection, activated CD8^+^ αβ T cells exhibited upregulation of pathways related to the transport of organic and nitrogen-containing compounds (NES= 1.46 to 1.43), and intracellular trafficking (NES= 1.58), reflecting enhanced cellular organization to support their effector functions. Biosynthetic pathways, including macromolecule and cellular biosynthesis (NES= 1.59), were also enriched, indicating increased metabolic and protein handling activity. In addition, pathways associated with cytokine production (NES= 1.79) (upregulation of *FCER1G*, *HSP90AA1*, *IRF1*) and epithelial development (NES= 1.83) were activated, consistent with their role in coordinating immune responses and controlling infection.

#### Cycling CD4^+^ and CD8^+^ αβ T cells

We found activation of carbohydrate metabolism (NES= 1.56), including genes activating glycolysis and gluconeogenesis, glycogen synthesis and metabolism and Krebs cycle. Phosphate metabolic process was also upregulated (NES= 1.44). Also, we observed that several immune signaling pathways including type I interferon production (NES= 1.85; upregulation of *IRF1*, *IFI6*), toll-like receptor signaling (NES= 1.86; upregulation of *CXCL10*, *FOS*, *CGAS*), and pattern recognition receptor pathways were upregulated. In contrast, processes related to cell cycle progression, chromosome segregation, and DNA replication were downregulated (NES = −2.4 to −1.6). Similarly, cytoskeleton organization was downregulated (NES= - 1.55).

#### Follicular CD4^+^ αβ T cells

Pathways involved in cytoplasmic translation, peptide and amide biosynthesis, ribosome biogenesis, and ribonucleoprotein complex assembly were downregulated (NES= −2.4 to −1.68), reflecting a reduction in global protein synthesis.

#### Non-naive CD4^+^ αβ T cells

Non-naive CD4^+^ αβ T cells, which include memory and effector subsets, play a central role in orchestrating adaptive immune responses during infection. There was a profound downregulation of genes involved in T cell differentiation and activation (*RUNX2*, *RORA*, *PTPN22*, *IL7R*, *CBLB, EGR1*) and regulation of cytokine production (*HMGB2*). On the other hand, we found upregulation of inflammatory genes such as *FOS*, *JUN*, *CXCL10*, as well as *POU2F2*, involved in glycolysis, was also upregulated.

#### IFN-responsive CD8^+^ T cells

Leukocyte differentiation genes such as *PTK2B*, *RORA*, *PTPN22*, and *GPR183* were downregulated, meanwhile other such as *TBC1D10C* (inhibitor of Ras signaling pathway) and *GIMAP1* were upregulated in this cell subset. Inflammatory genes *TAP1*, *IFI6, IRF1*, *CXCL10* and *JUNB* were downregulated, but genes coding for TFG-β and IL-12 receptors (*TGFBR2* and *IL12RB2*, respectively) were downregulated.

#### NK cells

NK cells CD8⁻ are innate cytotoxic lymphocytes that contribute to early control of infection by directly killing infected cells and producing cytokines that shape the immune response. During *S.* Typhimurium infection, these cells downregulated genes that led to inhibition (NES= −2.18) of mitochondrial respiration and energy production (complex I) and oxidative phosphorylation (*ATP5F1B*, *ATP5MC3*, *ATP5MG*, *ND4*, *NDUFB7*, *NDUFB9*, *COX1*, *COX2*, *COX3*). Pathways related to regulation of lymphocyte differentiation (NES= 1.85; *IFNG*, *IRF1*) and antigen processing (*TAP1*, *PDIA3*, *HSP90AA1*, *HSPA6*,) were also enriched, reflecting their role in coordinating immune responses.

CD8^+^ NK cells are a rarer subset of NK cells that have greater cytotoxicity capacity and stronger response to cytokine stimulation than the CD8^−^ NK subset. In porcine ileum we found that during *S.* Typhimurium infection these cells have a decreased regulation of chemotaxis (NES= −2.12), mainly due to downregulation of genes such as *CXCR4* and *DUSP5*.

## DISCUSSION

Controlling *S.* Typhimurium in swine production remains a major challenge, as this pathogen continues to circulate widely in pig farms and significantly contributes to human non-typhoidal salmonellosis (2, 22). Its ability to efficiently colonize the porcine intestine and persist within host tissues, even in the absence of overt clinical signs of disease, further complicates eradication efforts and facilitates chronic carriage, thereby enhancing transmission along the food chain (4, 23). Understanding the detailed pathogenesis of ileal infection, including the identity and functional state of specific immune and epithelial cell populations, is critical for both improving animal health and mitigating zoonotic risk. In this study, a complex and heterogeneous cellular landscape was revealed in ileum, comprising enterocytes, myeloid cells (monocytes, macrophages, and dendritic cells), innate lymphoid cells (ILCs), B cells, and diverse T cell subsets. These findings are consistent with previous studies describing immune populations in the porcine ileum and other intestinal regions, which report a predominance of T and B lymphocytes alongside substantial myeloid compartments (24, 25). Importantly, the high-resolution single-cell approach enabled the identification of fifteen T and ILC subtypes and five B cell subtypes, in addition to enterocytes and myeloid populations, including monocytes/macrophages and dendritic cells, thereby providing an unprecedented level of resolution within the infected ileum. Comparisons with human intestinal studies indicate a conserved architecture of immune populations between pigs and humans, reinforcing the translational relevance of the porcine model for enteric infections (26, 27). Recent porcine single-cell RNA-seq atlases of the pig intestine provide complementary evidence of intestinal cellular diversity, highlighting the validity and novelty of our single-cell observations in the context of infection (28). Finally, the detection of *Salmonella* associated transcriptional programs within specific cell types highlights the dynamic, cell type-specific response to infection, emphasizing mechanisms that may contribute to bacterial persistence or immune evasion.

A profound remodeling of the ileal immune landscape during *Salmonella* infection was found, characterized by coordinated shifts across B cells, T cells, myeloid populations, and the intestinal epithelium. B cell subsets displayed a heterogeneous but well-structured response: number of activated and cycling B cells expanded markedly, whereas resting, transitioning, and antibody-secreting B cells declined. This pattern is reminiscent of mucosal B-cell activation and differentiation dynamics recently described in porcine ileal and jejunal Peyeŕs patches under homeostatic conditions (25), suggesting that infection rapidly triggers an activation program that can override baseline differentiation states. Similarly, T-cell compartments underwent dynamic reshaping, with increased abundance of naïve CD4^+^ and CD8^+^ T cells, follicular CD4^+^ T cells, and NK T cells alongside reductions in activated, cytotoxic, and non-naïve CD8^+^ T cell as well as γδ T-cell subsets. These shifts reflect both systemic and local T-cell responses, highlighting the conserved recruitment and activation pathways in pigs following *Salmonella* infection (29).

Innate myeloid populations including monocytes, macrophages, and dendritic cells were significantly expanded in infected ileum, reflecting rapid recruitment in response to pathogen associated molecular patterns (PAMPs), including conserved microbial motifs such as LPS or flagellin, which trigger innate pattern-recognition receptors and promote inflammatory activation (30, 31). In contrast, group 3 ILCs and enterocytes decreased in abundance, consistent with epithelial injury, barrier disruption, and impaired mucosal homeostasis. These changes align with transcriptional signatures of inflammation and epithelial dysfunction reported in porcine ileum during *S*. Typhimurium infection (32).

The GSEA revealed pronounced cell type-specific transcriptional responses to *Salmonella* infection, reflecting both conserved and lineage-restricted pathways. Our data highlights that epithelial and immune compartments engage distinct yet complementary programs of inflammation, stress response, and immune modulation, revealing the complex orchestration of host defense at the single-cell level. Comparisons between infected and control samples identified pathways that were commonly affected across multiple cell types, as well as those selectively enriched or suppressed in specific lineages, suggesting that *Salmonella* elicits both broad and highly targeted host responses. Our data suggests that porcine enterocytes actively reprogram their metabolism, down-shifting pathways involved in lipid and amino-acid catabolism to prioritize immune and stress-related functions. This metabolic adaptation is consistent with previous work showing that *Salmonella* can alter epithelial metabolism and oxygenation, thereby modifying the luminal environment, which could affect colonization resistance (33). Suppression of lipid-catabolic genes may reflect a systemic shift toward glycolysis and away from the TCA cycle, a strategy observed in *Salmonella* infection to sustain rapid inflammatory responses (34). Activation of stress, ion-transport, and epithelial extrusion pathways likely represent a defensive response, consistent with inflammasome-mediated expulsion of infected cells, and enhanced mucin production aimed at maintaining epithelial integrity (35).

During *Salmonella* infection, porcine monocytes/macrophages, which are key phagocytic cells, undergo profound metabolic and immune reprogramming, characterized by enhanced glycolysis and activation of inflammatory and stress-response pathways. *S*. Typhimurium has been shown to upregulate glycolysis in macrophages as a strategy for intracellular survival (36). Other studies indicate that *Salmonella* can impair macrophage defense by disrupting glycolysis-dependent phagosome acidification, thereby undermining bacterial clearance (37). At the same time, dendritic cells, essential antigen-presenting cells that bridge innate and adaptive immunity and contribute to intestinal immune surveillance, likely mount a complex response (38). Although they activate inflammatory and interferon-mediated pathways, virulent *S*. Typhimurium strains can impair dendritic cell maturation and antigen presentation reducing their ability to effectively prime adaptive immunity and may also modulate dendritic cell function through interactions with CD209 receptors to facilitate dissemination and alter antigen-presenting capacity (39, 40).

In the porcine ileum during *S*. Typhimurium infection, ILCs show stress-response and antigen-presentation signatures. Group 3 ILCs upregulate heat-shock protein genes and TNF/MAPK pathway components, while group 1 ILCs increase MHC class I/II and IFN-γ related gene expression, reflecting a role in bridging innate sensing and adaptive T-cell activation. These findings are consistent with reports that intestinal ILCs adjust cytokine production in response to bacterial infection and can migrate via lymph to mesenteric lymph nodes while producing IFN-γ (41, 42). Overall, ILCs in the infected ileum integrate microbial and stress signals to contribute to early mucosal defense and coordinate adaptive immune responses.

In porcine ileum, CD8^−^ NK cells downregulate genes associated with mitochondrial respiration and oxidative phosphorylation, while upregulating lymphocyte differentiation and antigen-processing pathways. This suggests an early-response activation program in which these NK cells prioritize rapid cytokine production and immune coordination over energy-consuming mitochondrial metabolism. Such metabolic remodeling is consistent with the concept of metabolic flexibility, allowing NK cells to sustain effector functions under conditions of restricted mitochondrial activity (43, 44). The CD8^+^ NK subset, which is less abundant but more cytotoxic, shows downregulation of chemotaxis and cytoskeleton related genes such as *CXCR4* and *PFN1*. The downregulation of the former could reflect an altered migratory capacity or modified homing program (45–47).

B cells, particularly those residing in ileal Peyeŕs patches, play a central role in mucosal immunity, potentially contributing to IgA production and antigen sampling during enteric infection (48–50). During *Salmonella* infection, the different B-cell subtypes identified in our dataset reflect this functional specialization. The marked reduction of chemotactic, translational, and biosynthetic programs in activated and cycling B cells suggests a metabolic restriction that may help balance activation with energy conservation, especially under inflammatory stress (51). In contrast, antibody-secreting B cells exhibit strong induction of protein-folding and chaperone pathways together with increased oxidative phosphorylation, consistent with the reliance of these cells on mitochondrial respiration to support high-rate antibody secretion (52). The upregulation of antigen presentation genes in these B cells is compatible with B cells in Peyeŕs patches acting as antigen-presentation cells, rather than being limited to antibody production (53). Transitioning B cells, which show increased innate and interferon-related signatures, may reflect early microbial sensing in Peyeŕs patches prior to full differentiation. Resting B cells maintain mitochondrial organization and protein-folding machinery while reducing global protein synthesis, reflecting a quiescent state that preserves cellular homeostasis and readiness for activation. These dynamics are consistent with observations that resting B cells migrate into follicles to maintain homeostasis and prepare for rapid functional responses, as reported in porcine jejunal and ileal Peyeŕs patches (25).

T cells play a crucial role in controlling *S*. Typhimurium infection, orchestrating both adaptive and innate-like responses in the gut mucosa. Conventional αβ T cells such as CD4^+^, CD8^+^, naïve and memory subsets are adaptive lymphocytes that recognize antigens via the T-cell receptor (TCR) and require activation by antigen-presenting cells, mediating specific and long-lasting immunity through IFN-γ production and cytotoxic activity (54). In contrast, γδ T cells possess a hybrid phenotype, combining innate-like rapid responses with adaptive features: they recognize stress-induced or non-peptide antigens independently of classical MHC, act as early sentinels in the intestinal epithelium, and bridge innate and adaptive immunity (55). Understanding the transcriptional and functional heterogeneity of these T-cell subsets is therefore critical to elucidate host defense mechanism against enteric *Salmonella*. Our single-cell transcriptomic analysis reveals coordinated metabolic and stress-adaptive responses across T-cell subsets during infection. Activated CD8^+^ αβ T cells upregulate biosynthetic, trafficking, and cytokine production pathways, supporting effector functions such as IFN-γ secretion for bacterial clearance. Cycling CD4^+^ and CD8^+^ αβ T cells show enhanced glycolysis, carbohydrate metabolism, type I interferon and TLR signaling, balancing proliferation with immune readiness (56). Non-naïve CD4^+^ αβ T cells and IFN-responsive CD8^+^ T cells upregulate chaperone-mediated protein folding and stress-response pathways, maintaining function during sustained activation (56). Follicular CD4^+^ αβ T cells show increased protein folding but reduced global protein synthesis, consistent with germinal center support and B-cell help, while naïve CD4^+^ and CD8^+^ αβ T cells exhibit pre-activation stress signatures (57, 58). Collectively, these results indicate that T cells undergo metabolic and proteostatic remodeling, integrating rapid effector activity with long-term fitness while γδ T cells serve as early sentinels bridging innate and adaptive immunity in the gut mucosa.

Together, these findings provide a comprehensive single-cell map of the porcine ileal response to *S*. Typhimurium, revealing coordinated remodeling across epithelial, myeloid, B-cell, ILC, and T-cell compartments. The infections induce profound metabolic and stress-adaptation programs that shape both innate and adaptive immunity, while uncovering distinct functional specializations within each cellular lineage. By resolving heterogeneous immune and epithelial states that cannot be captured by bulk analysis, this work establishes a high-resolution framework for understanding host-pathogen interactions in the porcine gut and underscores the translational relevance of the pig as a model for human enteric salmonellosis, providing a strong foundation for future efforts to improve mucosal immunity, disease control, and vaccine development.

## MATERIALS AND METHODS

### Bacterial strain, experimental infection and sample collection

The *Salmonella enterica* subspecies *enterica* serovar Typhimurium phagetype DT104 SP11 was isolated from a carrier pig (59). Bacteria were grown at 37°C in LB broth to logarithmic stationary phase (OD_600_ = 0.8). Six weaned male and female crossbreed piglets, approximately 4 weeks of age, were used in this study. Prior to the experiment, all animalś fecal samples were confirmed to be free of *Salmonella*. Experimental procedures were performed at University of León (Spain) following a methodology previously described (32). Briefly, piglets were divided into two groups: three pigs were orally inoculated with 6.25 mL of sterile LB broth, (control group, C) and three pigs were orally inoculated with 6.25 mL of LB broth containing 5 × 10^8^ colony-forming units (CFU) of *S*. Typhimurium (*Salmonella* group, S). On day 2 (acute infection), piglets were humanely euthanized, necropsied, and ileum tissue samples were collected. After resection, the ileal contents were gently squeezed out, the lumen was washed with sterile PBS, and the ileum was cut into fraction of around 10 cm and were immediately placed in Hypothermosol® for subsequent cell dissociation. Tissues collected for histological processing were immediately fixed in 4% paraformaldehyde in 0.1 M phosphate buffer (pH 7.4). Symptoms of infection, including fever, diarrhea, and lethargy, were recorded daily for each animal to monitor disease progression. All animal handling and experimental procedures complied with the Good Experimental Practices (GEP) guidelines and were approved by the Ethical and Animal Welfare Committee of the University of León (Spain).

### Detection and Quantification of Salmonella in Feces

Feces from pigs were tested by qPCR for *Salmonella* to ensure the effectiveness of the infected challenge. DNA was extracted from 200 mg of frozen fecal samples using the PureLink^TM^ Microbiome DNA Purification Kit (Invitrogen, Thermo Fisher Scientific Inc., Waltham, MA, USA) following the manufacturer’s instructions. Quantification of *S.* Typhimurium was performed by TaqMan quantitative polymerase chain reaction (qPCR) using a previously described assay (60). To generate the standard curve, DNA from a pure culture of the same *Salmonella* strain used in the challenge was extracted with the same kit and serially diluted to known genome equivalents (1.0 ×10^4^, 5.0 ×10^3^, 1.0 ×10^3^, 5.0 ×10^2^, 2.5 ×10^2^, 1.0 ×10^2^,5.0 ×10^1^ GEd per μl, plus 0 for negative control), with 1 genome equivalent corresponding to 5.46904 fg of DNA. The qPCR reaction employed a 19-mer forward primer (5′-GCGCACCTCAACATCTTTC-3′), a 21-mer reverse primer (5′-CGGTCAAATAACCCACGTTCA-3′), and a FAM-labeled fluorogenic probe (FAM-ATCATCGTCGACATGC-MGB/NFQ). Reactions (25 μl) contained 12.5 μl IQ Supermix 2X (Biorad, Madrid, Spain), 0.4 μM of each primer, 0.2 μM probe, 1 μM MgCl₂, 200 ng DNA, and nuclease-free water. Amplifications were performed on an iQ5 Thermocycler (Biorad, Madrid, Spain) with the following cycling conditions: 95 °C for 3 min, followed by 50 cycles of 95 °C for 15 s and 60 °C for 1 min.

### Immunohistochemistry and immunofluorescence using a S. Typhimurium specific antibody

Tissue samples were embedded in paraffin, sectioned at 5 μm, and mounted on poly-L-lysine–coated slides (Sigma-Aldrich, St. Louis, MO, USA). Sections were dried overnight at 37 °C, deparaffinized in xylene, and rehydrated through graded alcohols to distilled water. For immunohistochemistry (IHC), antigen retrieval was performed by heat treatment in 0.01 M citrate buffer. Sections were incubated overnight at 4 °C with a rabbit polyclonal anti-*Salmonella* antibody (AgH, 1:200), followed by a 1 h incubation at 37 °C with a biotinylated anti-rabbit IgG secondary antibody (Dako, Barcelona, Spain). Antibody binding was visualized using a standard avidin–biotin peroxidase system (Dako, Barcelona, Spain) with diaminobenzidine (Sigma-Aldrich, St. Louis, MO, USA) as the chromogenic substrate. To assess specificity, negative controls omitting the primary antibody were included in each assay. After development, sections were counterstained with hematoxylin, dehydrated, and mounted.

For immunofluorescence (IF), serial sections were processed in parallel using the same primary antibody under identical incubation conditions. As a secondary antibody, Alexa Fluor 594 anti-rabbit IgG (Life Technologies, Carlsbad, CA, USA) was used and incubated for 1 h at 37 °C. Slides were washed three times for 5 min each in PBS containing 1.43 μM 4′,6-diamidino-2-phenylindole (DAPI; Life Technologies) for nuclear counterstaining. Fluorescent signals were visualized using a Thunder microscope (Leica Microsystems), and representative images were analyzed to assess the localization of *Salmonella* within the intestinal tissue.

### Single cell RNA sequencing (scRNA-seq)

The tissue was incubated in mucus dissociation buffer (Hank’s Balanced Salt Solution (HBSS, Sigma-Aldrich) + 5 mM dithiothreitol (DTT) + 2% Fetal Bovine Serum (FBS)) for 20 min on ice, washed with 1x PBS and the serosa was peeled off and minced finely. BLP digestion buffer (2 mM EDTA in HBSS+BLP (260 U/20 mg)) was added and, after vigorous shaking every 30 s for 10 min on ice, Col I/IV+DNase I digestion buffer (HBSS+5 mg/mL collagenase I+2.5 mg/mL collagenase IV+15 μg/mL DNase I) was added for 30 min at room temperature. The suspension was then diluted with a cold HBSS buffer and passed first through a 100-μm strainer and then through a 40-μm strainer. It was then centrifuged at 600 x g for 5 min and 1 mL of red blood cell (RBC) lysis buffer was added for 3 min at room temperature. Finally, two washes were performed with ice-cold wash buffer (1% BSA in HBSS), and the cells were counted using trypan blue in an automatic cell counter (Biorad, Madrid, Spain). A total of 1.5 × 10^6^ cells were transferred to a microtube, washed twice with 1x DPBS buffer (Dulbeccós Phosphate Buffered Saline), and resuspended in 300 μL of 1x DPBS. The suspension was transferred to a cryovial, and four volumes of chilled 100% methanol were added dropwise while gently mixing to prevent clumping. Cells were incubated for 30 min at −20°C and subsequently shipped to the National Center for Genomic Analysis (CNAG, Barcelona) for scRNA-seq analysis. At CNAG, the cryovial was equilibrated on ice for 5 min, followed by centrifugation and removal of the methanol supernatant. Cells were resuspended in Wash-Resuspension buffer to a final concentration of 1,000 cells/μL and immediately processed using the 10x Genomics Single Cell Gene Expression protocol.

Single-cell suspensions were processed using the Chromium Single Cell platform (10x Genomics, USA) following the manufactureŕs protocol for single-cell 3’ gene expression library preparation. Briefly, single cells were encapsulated into Gel Bead-in-Emulsions (GEMs) to enable barcoding of individual transcripts. Complementary DNA synthesis, amplification, and library construction were performed according to the manufactureŕs recommendations. The resulting libraries were sequenced on an Illumina high-throughput sequencing platform using paired-end sequencing.

### Bioinformatic data analysis

We generated the *Sus scrofa* reference genome (primary assembly from Ensembl https://ftp.ensembl.org/pub/release-112/fasta/sus_scrofa/dna/Sus_scrofa.Sscrofa11.1.dna.toplevel.fa.gz) and transcriptome (https://ftp.ensembl.org/pub/release-112/gtf/sus_scrofa/Sus_scrofa.Sscrofa11.1.112.gtf.gz) using the official 10x Genomics instructions for custom reference construction. Raw sequencing data were processed with Cell Ranger v7.0, which performed read alignment, UMI quantification, and generation of the gene-by-cell expression matrix.

First, the CellBender (v0.3.0) software, with parameters –fpr 0.01 and –epochs 150, was used to correct for the presence of ambient RNA, which is the RNA that is outside the cells after cell death and is present in the empty droplets (61). Quality control filtering was performed to exclude low-quality or aberrant barcodes. We first assessed library size, and required cells to contain at least 750 UMIs, as barcodes with fewer counts typically represent empty droplets or ambient contamination. Similarly, cells with fewer than 200 detected genes were removed, as these profiles likely correspond to droplets lacking true cellular material. Mitochondrial read fraction was also used as an indicator of cell integrity: cells with >6% mitochondrial UMIs were considered low quality, as elevated mitochondrial content often reflects damaged or dying cells that have lost cytoplasmic RNA while retaining mitochondrial transcripts. To further refine QC, we examined the distribution of intronic content using DropletQC v0.0.0.9000 (62, 63). Barcodes with <8% intronic UMIs were removed, as these likely represent empty droplets dominated by ambient RNA, whereas droplets with >62% intronic UMIs were excluded because they tend to correspond to damaged or nuclei-like profiles. At the upper end of the distribution, barcodes with >50,000 UMIs or >6,000 detected genes were also filtered out, as these unusually large profiles frequently reflect multiple or technical artifacts.

After this initial QC filtering, we performed doublet prediction using DoubletFinder v2.0.4 (64), (estimating parameters following the recommendations in its documentation and removed all predicted doublets). Final downstream analyses—including normalization, dimensionality reduction, clustering, and visualization—were carried out using Seurat v5.1.0 (65). For downstream preprocessing, genes with zero counts across all cells and genes expressed in fewer than five cells were removed. Highly variable features were identified using the FindVariableFeatures() function, selecting the top 3,000 most variable genes, and expression values were subsequently scaled and centered using ScaleData(). Dimensionality reduction was performed using principal component analysis (RunPCA()). To correct for batch effects, samples were integrated using Harmony (RunHarmony()) on the first 30 principal components. Clusters were identified by constructing a shared nearest-neighbor graph with FindNeighbors() using these 30 Harmony dimensions, followed by community detection with FindClusters() at a resolution of 0.8. Finally, UMAP embeddings were generated using the same 30 Harmony dimensions via RunUMAP().

Cell type annotation was performed primarily by label transfer using a published porcine ileum scRNA-seq reference dataset (24). This reference contains 26 annotated cell types spanning major immune and non-immune lineages, and closely matches the tissues profiled in the present study. Genes shared between the reference and query datasets were identified and used to compute integration anchors with FindTransferAnchors() (PCA reduction using 30 dimensions). Cell type labels were then assigned using TransferData(), which generates a prediction score for each reference cell type for every cell in the query dataset. Each cell was annotated with the label corresponding to its highest prediction score, retaining only assignments with a confidence score >0.50 and labeling the remainder as “unknown.” Cluster-level labels were derived by determining the predominant cell type within each cluster and assigning that label when it represented ≥25% of cells in that cluster. The resulting annotations were used to characterize the distribution of cell populations in the porcine ileum.

To evaluate changes in cell type composition between conditions, we performed a differential abundance analysis based on contingency testing. For each annotated cell type, we generated a cell-by-sample count matrix and aggregated cell numbers across biological replicates belonging to the Control or Infected groups. For each cell type, a 2×2 contingency table was constructed comparing (i) the number of cells of that cell type in Control versus Infected samples, and (ii) the number of all remaining cell types in Control versus Infected samples. A Fisher’s exact test was applied to each contingency table to assess whether the proportion of that cell type differed significantly between groups. Resulting p-values were corrected for multiple testing using the Benjamini– Hochberg false discovery rate (FDR) procedure. Differentially abundant cell types were identified based on their FDR-adjusted p-values. The study was carried out for each annotated cell subtype by label transfer. Using FindMarkers, differentially expressed genes (DEGs) were found between controls and infected using the Wilcoxon test. Results were filtered keeping only those with an adjusted p-value < 0.05 for future analysis.

### Functional enrichment analysis

To identify the signaling pathways and biological processes modulated in response to intestinal *Salmonella* infection, functional enrichment analyses were performed using complementary approaches. DEGs from each cell type identified in the porcine ileum were used as input for downstream analyses.

Gene Set Enrichment Analysis (GSEA) was applied to detect coordinated gene expression patterns within functional gene sets. The Hallmark gene set collection from the Molecular Signatures Database (MSigDB; *h.all.v2024.1.Hs.symbols.gmt*) was used, as it summarizes well-characterized and minimally overlapping biological processed. GSEA results were interpreted using the Normalized Enrichment Score (NES). This approach enabled the identification of major functional pathways either activated or repressed across distinct cellular lineages in response to infection, providing an integrative view of the immunological, inflammatory, and metabolic responses at the single-cell level. In parallel, ontology-based functional enrichment was performed using Metascapés standard workflow (21). DEGs from each cell population were analyzed across multiple pathway and ontology databases integrated by Metascape, including GO Biological Process, KEGG Pathway, and Reactome. Enrichment statistics were calculated using the default hypergeometric test with Benjamini-Hochberg correction. Terms passing Metascape significance thresholds were further clustered based on Kappa similarity (>0.3) to generate non-redundant functional modules. Representative terms from each cluster were selected, and a subset of enriched terms was retained according to Metascapés recommended limits to generate enrichment networks and visualize relationships between functional categories. Network graphs were constructed by linking term pairs whose similarity exceeded the 0.3 threshold, and up to 15 terms per cluster and a maximum of 250 terms were included in network visualizations, with node attributes reflecting significance and gene-set membership.

## DATA AVAILABILITY

Data will be made available by request upon publication.

## AUTHORS CONTRIBUTIONS

Conceptualization and funding acquisition: S.Z.L. and J.J.G. Experimental work: J.M.S.C., R.F.R., A.R.G., T.G.G., C.A., A.M.M., J.J.G. and S.Z.L. Data analysis: J.M.S.C., T.M.A., M.A.N., A.E.C. and S.Z.L. Writing original draft: J.M.S.C., S.Z.L. Manuscript draft revisions and editing: A.E.C., T. M. A. Review and approval of draft: all authors.

## ACKNOWLEDGEMENTS

The authors wish to express their gratitude to Héctor Arguello and Ana Carvajal (Department of Animal Health, Faculty of Veterinary Medicine, University of León, Spain) for their valuable support during the *in vivo* infection experiments.

